# Beyond thermal unfolding: urea-gradient nanoDSF approach for thermostability analysis of kinetically stable hyperthermophilic proteins

**DOI:** 10.64898/2026.04.09.717478

**Authors:** Wojciech Rusinek, Sebastian Dorawa

**Author notes:** Correspondence: S. Dorawa, Laboratory of Extremophiles Biology, Department of Microbiology, Faculty of Biology, University of Gdansk, Wita Stwosza 59, 80-308 Gdansk, Poland.

## Abstract

In this study, we demonstrate that urea enables reliable melting temperature (T_m_) determination of hyperthermostable proteins by nano differential scanning fluorimetry (nanoDSF) Under native conditions, Pfu DNA polymerase and its Sso7d-fusion variant showed no detectable unfolding transitions, despite their T_m_ values falling within the instrument’s operational range, reflecting their extreme kinetic stability. In the presence of up to 7 M urea, intrinsic tyrosine and tryptophan fluorescence revealed clear unfolding transitions, yielding extrapolated T_m_ values of 104.8 ± 0.09 °C for Pfu and 106.8 ± 0.33 °C for its Sso7d-fusion variant. These results demonstrate that urea-gradient nanoDSF overcomes both instrumental and kinetic limitations, providing a simple and robust method for assessing the thermal stability of (hyper)thermostable proteins.

**Graphical abstract:** 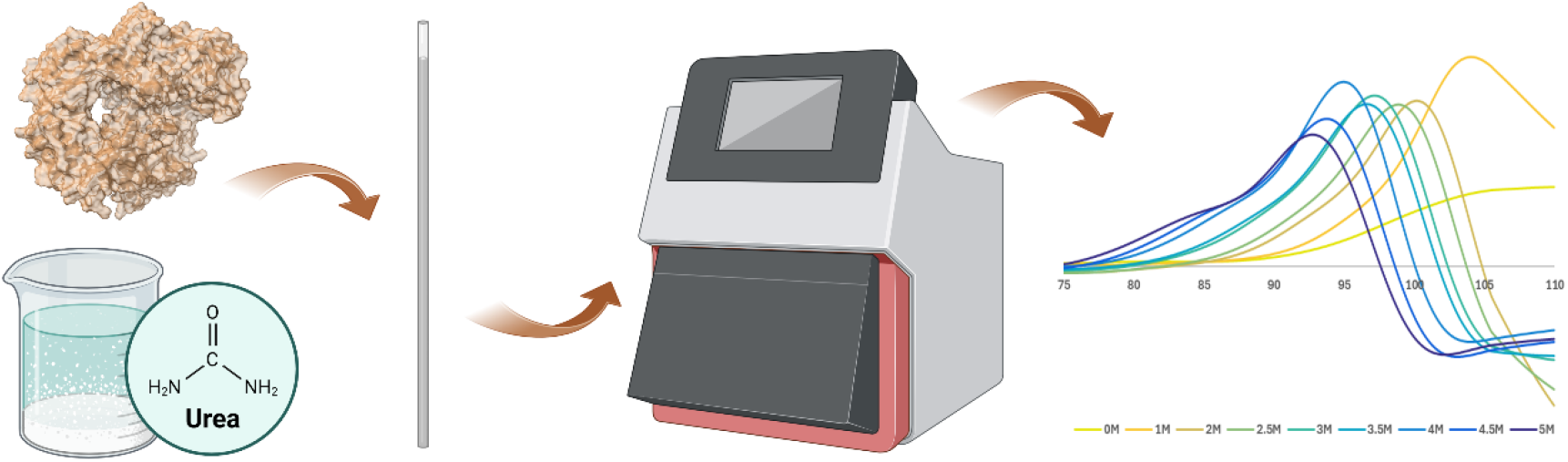

## 1. Introduction

Proteins derived from thermophilic (growth temperature ∼45-75 °C) and hyperthermophilic (>80 °C) organisms are characterized by exceptional thermal stability and resistance to chemical denaturants [1,2]. These properties make them highly attractive for biotechnological applications, but at the same time pose significant challenges for their biophysical characterization [3,4]. In particular, thermal unfolding of such proteins often occurs at temperatures exceeding the operational limits of commonly used analytical techniques.

Thermal stability is a fundamental property of proteins, defining the range over which they maintain their native structure and biological activity [5,6]. As protein function is intrinsically linked to structural integrity, there is a clear need for robust methods enabling the analysis of protein folding and stability under a wide range of conditions.

Conventional approaches for protein thermal stability assessment, such as differential scanning calorimetry (DSC) and circular dichroism (CD), are instrumentally demanding, require relatively large amounts of protein, and offer limited throughput [7]. Moreover, their applicable temperature ranges often preclude the analysis of highly thermostable proteins, highlighting the need for alternative approaches better suited to such systems. Nano differential scanning fluorimetry (nanoDSF) addresses many of these limitations by monitoring intrinsic tryptophan and tyrosine fluorescence in a label-free, high-throughput format with minimal sample consumption [8]. This approach has been successfully applied to the analysis of proteins from extremophilic environments, including thermostable bacteriophage-derived enzymes [9,10]. However, even nanoDSF instruments with an upper temperature limit of 110 °C may fail to capture unfolding transitions of some hyperthermostable proteins [11].

To overcome these limitations, chemical denaturants can be used to modulate protein stability and shift unfolding transitions to lower, experimentally accessible temperatures [12]. Urea was selected as the chemical denaturant due to its milder and more gradual destabilizing effect compared to guanidine hydrochloride [13], to ensure more precise detection of thermal unfolding transitions. Here, we establish a urea-gradient nanoDSF approach as a robust and broadly applicable strategy for probing the thermal stability of hyperthermostable proteins. Using two widely utilized model systems: Pfu DNA polymerase and its Sso7d-fusion variant, we demonstrate that this method enables accurate and reproducible melting temperature determination under conditions fully compatible with the Prometheus NT.48 platform, thereby extending the applicability of nanoDSF to proteins previously inaccessible to reliable thermal analysis.

## Materials and Methods

### Bacterial Strains, Plasmid, and Materials

The *E. coli* Rosetta 2(DE3)pLysS (Merck KGaA, Darmstadt, Germany; cat. no. 71403-3) was used for protein overproduction. Cells were grown in Lysogenic broth (LB) or on Lysogenic agar (LA) plates [13]. When required, media were supplemented with chloramphenicol (34 µg/mL) or kanamycin (30 µg/mL). Protein overexpression was carried out using the expression vectors pET24a(+)_Pfu and pET24a(+)_Pfu-Sso7d.

### Expression and Purification of Recombinant Proteins

Pfu and Pfu-Sso7d DNA polymerases were expressed in *E. coli* Rosetta 2(DE3)pLysS cells harboring the appropriate expression plasmids. The cultures were cultivated in LB medium supplemented with chloramphenicol and kanamycin at 37 °C until reaching an OD_600_ of 0.5. Protein expression was then induced by the addition of 1 mM IPTG (Serva, Heidelberg, Germany; cat. no. 26600.06), followed by incubation for 4 h at 37 °C. Cells were harvested by centrifugation (5,000 × g, 20 min, 4 °C) and stored at −80 °C.

Cell pellets were resuspended in buffer A (50 mM NaH_2_PO_4_, pH 8.0, 500 mM NaCl, 0.1% [v/v] Triton X-100, 5% [v/v] glycerol and 10 mM imidazole) and disrupted by sonication on ice (30 cycles of 10 s at an amplitude of 12 μm) using a MISONIX XL2020 sonicator (Misonix Inc., Farmingdale, NY, USA). Cell debris was removed by centrifugation (10,000 × g, 20 min, 4 °C). The supernatant was incubated at 75 °C for 30 min to denature thermolabile host proteins, followed by a second centrifugation step (10,000 × g, 20 min, 4 °C) to obtain clarified lysate. Clarified lysate was applied to a 1 mL HiTrap TALON Crude column (Cytiva, Uppsala, Sweden; cat. no. 28-9537-66) connected to an ÄKTA pure 25 chromatography system (Cytiva, Uppsala, Sweden). The column was washed with 10 mL of buffer A, followed by an additional wash with buffer B (buffer A supplemented with 20 mM imidazole). Bound proteins were subsequently eluted with buffer C (buffer A supplemented with 200 mM imidazole). Peak fractions containing substantial quantities of the target protein were pooled and dialyzed against buffer D (20 mM Tris-HCl, pH 8.0, 50 mM NaCl and 5% [v/v] glycerol). For further purification, DNA polymerases were loaded onto a 1 mL HiTrap Heparin HP column (Cytiva, Uppsala, Sweden; cat. no. 17-0406-01) equilibrated with buffer D. The column was washed with an equilibration buffer, and proteins were eluted using a linear gradient of 50-1,000 mM NaCl in buffer D. Fractions containing the target proteins were pooled and dialyzed against storage buffer (20 mM Tris-HCl, pH 7.4, 50 mM KCl, 1 mM DTT, 0.1 mM EDTA, 0.5% Tween-20, 0.5% Triton X-100 and 50% [v/v] glycerol), and stored at -20 °C. All purification steps were monitored by SDS–PAGE, and protein concentrations were determined using the Bradford assay [14].

### Thermostability Analysis

Thermal stability was analyzed by nanoDSF with a Prometheus NT.48 instrument (NanoTemper Technologies, Munich, Germany). Standard-grade glass capillaries (NanoTemper Technologies, Munich, Germany; cat. no. PR-C002) were filled with 10 μL of recombinant protein stock solution, supplemented with 90 μL of 0–7 M urea solution (Merck KGaA, Darmstadt, Germany; cat. no. U5378). Prior to analysis samples were left to equilibrate for 5 min at room temperature then were centrifuged at 4,000 × g for 5 min at room temperature. Samples were heated from 20 °C to 110 °C at a rate of 0.5

°C/min. Protein unfolding was monitored by intrinsic tryptophan and tyrosine fluorescence at emission wavelengths of 330 and 350 nm, respectively. Data were analyzed using PR.StabilityAnalysis software (NanoTemper Technologies, Munich, Germany). Melting temperatures were determined according to the manufacturer’s instructions. All measurements were performed in triplicate (n=3).

## Results and Discussion

Conventional techniques such as DSC and CD are commonly used to assess protein thermal stability, but are limited by temperature range, sample requirements, or throughput, restricting their applicability to hyperthermostable proteins [15,16]. While nanoDSF offers high sensitivity and low sample consumption, its practical application is sometimes constrained by the inability to detect unfolding transitions for highly stable proteins. For example, Pfu DNA polymerase and its Sso7d-fusion variant pose a real challenge for biophysical characterization because their T_m_ exceeds the detection limit [17]. Previous studies have employed chemical denaturants such as urea and guanidine hydrochloride in DSC and CD measurements [18,19]; however, to our knowledge, this is the first application of a urea gradient to enable T_m_determination using nanoDSF.

We obtained highly pure recombinant proteins via overexpression followed by three-step purification protocol. SDS-PAGE analysis confirmed the homogeneity of the samples used in subsequent experiments (Fig. 1). The purification was efficient, yielding approximately 5 mg of Pfu DNA polymerase and 1.8 mg of Sso7d-fusion variant per liter of culture. Both proteins were enzymatically active, as confirmed by PCR-based assays (data not shown).

**Fig. 1.**
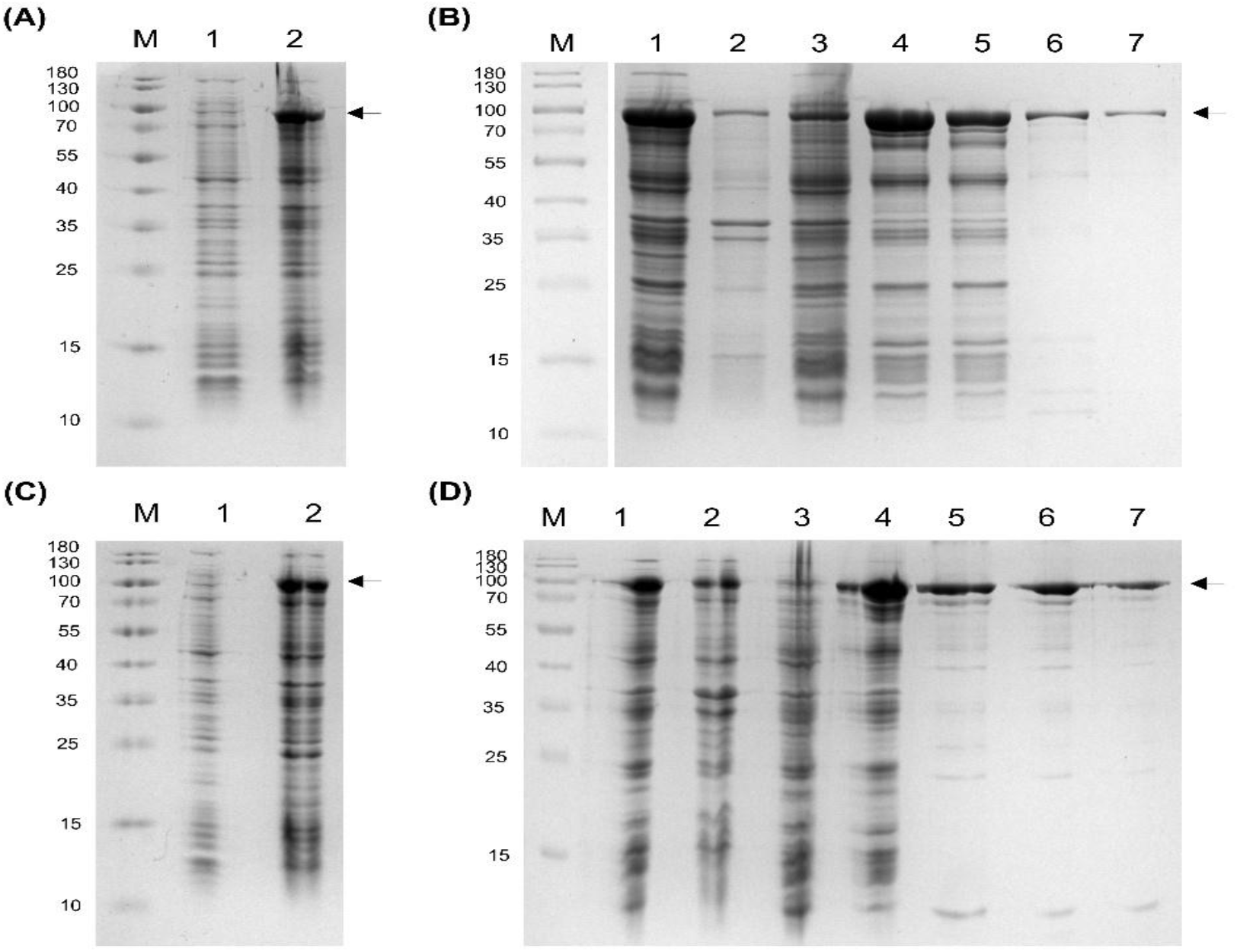
Overproduction and purification of Pfu DNA polymerase and its Sso7d-fusion variant. (A) Overproduction of Pfu DNA polymerase in *E. coli* Rosetta 2(DE3)[pET24a(+)_Pfu]. Lane M, molecular weight marker (10-180 kDa); lane 1, before induction; lane 2, after induction with 1 mM IPTG. (B) Purification of Pfu DNA polymerase. Lane M, molecular weight marker; lane 1, crude lysate; lane 2, insoluble fraction; lane 3, soluble fraction; lane 4, heat-treated lysate (75 °C, 30 min) after centrifugation (10,000 × g, 20 min, 4 °C); lanes 5 flow-through fractions from the HiTrap TALON crude column; lane 6, flow-through from the HiTrap Heparin HP column; lane 7, dialyzed protein. (C) Overproduction of the Pfu-Sso7d fusion protein in E. coli Rosetta 2(DE3)[pET24a(+)_Pfu- Sso7d]. Lane M, molecular weight marker; lane 1, before induction; lane 2, after induction with 1 mM IPTG. (D) Purification of the Pfu-Sso7d fusion protein under conditions analogous to (b). Proteins were separated by 12.5% SDS–PAGE and visualized by Coomassie Brilliant Blue staining. Arrows indicate the position of the His-tagged recombinant proteins.

Initial attempts to measure the thermal stability of the purified proteins under standard nanoDSF conditions failed to reveal detectable unfolding transitions. Proteins were subsequently analyzed in the presence of increasing urea concentrations (0−7 M), which allowed the detection of thermal unfolding transitions for both variants (Fig. 2A, B).

**Fig. 2.**
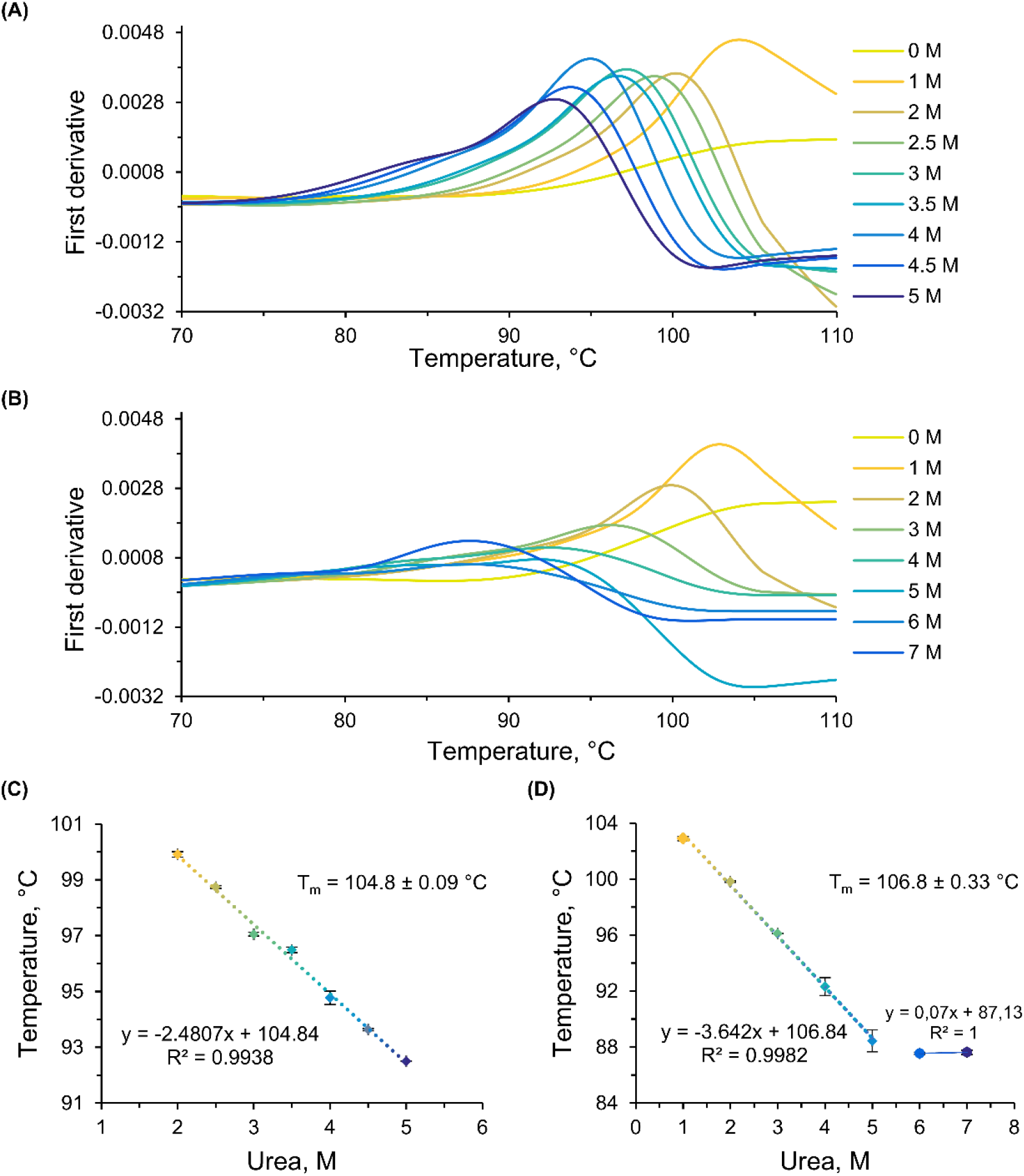
Thermal stability analysis of Pfu DNA polymerase and its Sso7d-fusion variant using urea-gradient nanoDSF. (A) Thermal unfolding of Pfu DNA polymerase monitored by nanoDSF in the presence of 0−5 M urea. (B) Thermal unfolding of the Pfu-Sso7d fusion protein monitored by nanoDSF in the presence of 0−7 M urea. The y-axis represents the first derivative of the fluorescence ratio (350/330 nm). Apparent T_m_ at each urea concentration were determined from the peak maxima using PR.StabilityAnalysis and are shown as mean ± SD (n = 3). (C,D) Linear dependence of T_m_ on urea concentration for Pfu DNA polymerase (2−5 M) and its Sso7d-fusion variant (1−5 M), respectively. Extrapolation to zero denaturant concentration was used to estimate the T_m_ in the absence of urea. Linear fits and corresponding coefficients of determination (R^2^) are shown.

For both proteins, T_m_ decreased linearly with increasing urea concentration (Fig. 2C, D). However, for the Sso7d-fusion variant, concentrations exceeding 5 M urea resulted in signal saturation (data points from 6-7 M), yielding an artifactual values R^2^ = 1.0 and T_m_ = 87.1 °C inconsistent with the linear trend observed at lower concentrations and therefore excluded from the extrapolation. Linear regression of the valid data range yielded extrapolated melting temperatures of 104.8 ± 0.09 °C for Pfu DNA polymerase and 106.8 ± 0.33 °C for its Sso7d-fusion variant, indicating a stabilizing effect of the fused Sso7d domain on the overall thermal stability of the enzyme.

A particularly noteworthy observation was the inability of the Prometheus NT.48 instrument to detect unfolding transitions for Pfu DNA polymerase and its Sso7d-fusion variant in the absence of urea, despite extrapolated T_m_ values (104.8 and 106.8 °C) falling within the nominal temperature range of the instrument (20-110 °C). This indicates that the limitation arises not solely from instrumental constraints, but rather from the intrinsic kinetic resistance of hyperthermostable proteins to unfolding. As previously reported, such proteins exhibit markedly slow unfolding rates, often several orders of magnitude lower than those of mesophilic counterparts, preventing equilibration on the timescale of standard thermal scans [21]. Consistent with this, no reliable unfolding transitions were observed even at a reduced heating rate of 0.1 °C/min (data not shown).

The addition of urea effectively overcomes this limitation by lowering both the thermodynamic stability and the kinetic barrier to unfolding [22], thus enabling the detection of well-defined unfolding transitions and reliable T_m_ determination.

## Conclusions

This work establishes urea-gradient nanoDSF as a powerful and broadly applicable approach for extending thermal unfolding analysis to hyperthermostable proteins, including those with extreme kinetic stability. Building on these findings, we propose a set of practical guidelines and a streamlined workflow for reliable melting temperature determination of hyperthermostable proteins using urea-gradient nanoDSF:

1. **Bioinformatic analysis** of the target protein sequence should be performed prior to any experimental work to confirm the presence of tryptophan and/or tyrosine residues, which are essential for intrinsic fluorescence-based detection in nanoDSF.
2. **The protein sample must be efficiently purified** and can be stored in the presence of glycerol as a cryoprotectant. Prior to measurement, the stock solution (e.g. 50% glycerol with protein concentration ≥ 1 mg/mL) should be appropriately diluted to reduce the **final glycerol concentration not exceeding 5%**, ensuring both sample stability during storage and proper signal quality during nanoDSF measurements.
3. **Urea-containing buffers** should be prepared fresh, best on the day of measurement to prevent carbamylation of protein residues resulting from urea decomposition [23].
4. **We recommend using a urea concentration range of 0−7 M** as a starting point, which can be further refined by adding intermediate values (e.g. 3.5, 4.5 M) if higher resolution of the linear range is needed. The upper boundary of the range must be carefully monitored. As observed for fusion variant examined in this study, concentrations exceeding 5 M resulted in signal saturation, rendering those data points unsuitable for extrapolation (Fig. 2D). Regardless of the range applied, **we strongly recommend restricting the analysis to the confirmed linear range only, while applying a strict R**^**2**^ **≥ 0.99 threshold as a quality criterion for linear regression**.
5. **Instrument parameters should be adjusted to the specific protein under investigation**. In our hands, a ramp rate of 0.5 °C/min over a temperature range of 20-110 °C, and laser excitation power of 70-100% respectively, yielded optimal unfolding curves for hyperthermostable Pfu and Pfu-Sso7d DNA polymerases.

## Abbreviations

nanoDSF: nano differential scanning fluorimetry
DSC: differential scanning calorimetry
CD: circular dichroism
T_m_: melting temperature.

## Acknowledgments

This work was supported by the University of Gdansk for financing through Research Projects of Young Scientists of the Faculty of Biology 2025 (grant no. 539-D100_B233_25) and National Science Centre, Poland, for financing under the MINIATURA 9 project (grant no. 2025/09/X/NZ1/00075). The equipment used in this study was funded by a capital equipment grant from the Polish Ministry of Education and Science (IA/SP/453344/2020), awarded to Dr. Anna-Karina Kaczorowska, supporting the Platform for Comprehensive Protein Characterization (Prometheus NT.48). The authors would like to thank the Collection of Plasmids and Microorganisms for providing access to research infrastructure, including the Prometheus NT.48 instrument. We are also grateful to Dr. Olesia Werbowy for kindly providing the expression plasmid carrying Pfu and Pfu-Sso7d DNA polymerases and for Dr. Aleksandra Kocot for valuable comments on the manuscript.

## Author contribution

**Wojciech Rusinek:** Conceptualization, Data curation, Formal analysis, Funding acquisition, Investigation, Methodology, Validation, Visualization, Writing – original draft. **Sebastian Dorawa:** Conceptualization, Formal analysis, Funding acquisition, Investigation, Methodology, Validation, Visualization, Supervision, Writing – original draft, Writing – review and editing

## Data accessibility

The data that support the findings of this study are available on request from the corresponding author.

